# Individual bounded rationality destabilizes cooperative dynamics in human–AI groups

**DOI:** 10.64898/2025.11.30.691459

**Authors:** Kazushi Tsutsui, Nobuaki Mizuguchi, Yuta Goto, Ryoji Onagawa, Fumihiro Kano, Kazutoshi Kudo, Tadao Isaka, Keisuke Fujii

**Affiliations:** Graduate School of Arts and Sciences, The University of Tokyo, Japan; Institute for Advanced Research, Nagoya University, Japan; Graduate School of Informatics, Nagoya University, Japan; Center for Advanced Intelligence Project, RIKEN, Japan; Research Organization of Science and Technology, Ritsumeikan University, Japan; Institute of Advanced Research for Sport and Health Science, Ritsumeikan University, Japan; Juntendo Administration for Sports, Health and Medical Sciences, Juntendo University, Japan; Faculty of Law, Meijo University, Japan; Department of Psychology, Queen’s University, Canada; Centre for the Advanced Study of Collective Behaviour, University of Konstanz, Germany; Department of Collective Behavior, Max Planck Institute of Animal Behavior, Germany; College of Sport and Health Science, Ritsumeikan University, Japan

## Abstract

Cooperative dynamics arise from interactions among individuals, including humans and artificial agents, yet it remains unclear how individual-level cognition and motion shape collective dynamics in such human–AI groups. We combined a dynamic cooperative task, in which one attacker was controlled by a human and two by heuristic artificial agents facing a deep reinforcement learning defender, with counterfactual simulations of artificial groups. Across nine task conditions, groups including a human consistently underperformed relative to groups composed solely of artificial agents. Humans exhibited restricted movement options and probabilistic rather than deterministic action choices, which shifted coordination dynamics towards less symmetric, less organized patterns and reduced performance for both individuals and groups. Embedding these boundedly rational movement and decision parameters into artificial agents reproduced human-like behavior and performance declines. These results demonstrate how limitations of a single individual can destabilize cooperative dynamics and provide insights into the design of robust human–AI systems.

## Introduction

Decisions are often made within social contexts^1,2^. Individuals are embedded within social groups that they both influence and are influenced by, and these reciprocal social influences shape the dynamics of collectives^3–5^. The collective intelligence that emerges from these interactions underpins not only the remarkable ecological success but also the destructiveness of the human species^6–8^. In modern society, a deeper understanding of these dynamic social processes seems more important than ever, both in human-to-human interactions and in our coexistence with other social entities such as artificial intelligence^9,10^.

Collective behavior is a complex, dynamic phenomenon that both emerges from and shapes individual cognition and motion^11,12^. Because of this inherent complexity, most previous studies have either abstracted away from individual sensorimotor processes and focused on macro-level dynamics^13–15^, or used constrained, simplified experiments to investigate specific aspects of decisions in social contexts^16–18^. Both approaches have been fruitful and have provided valuable insights into collective dynamics and decisions. Yet, focusing exclusively on one side of the dynamic interplay between individuals and collectives will necessarily overlook crucial aspects of social dynamics^19^. Few studies have quantified intertwined individual and collective dynamics within social groups, and the collective consequences of individual cognition and motion remain largely unexplored.

Here, to investigate how individual-level cognition and motion contribute to, or constrain, collective dynamics, we developed a novel triangular passing task in which humans and artificial agents could directly interact. This task represents a natural situation widely observed in real-world team ball sports^20–23^, while also providing a tractable framework for studying the link between individual decision-making and collective dynamics. In this task, three attackers attempted to maintain possession against a single defender. Notably, an individual decision made by one member of the attacking team could immediately affect the behavior of others, and those consequences could in turn feed back onto the individual. For example, when one attacker possessed the ball, other attackers might move to secure passing lanes for their teammate. However, such movements could distort the triangular formation, potentially narrowing the passing options available to themselves once they received the ball. Thus, while coordinated movement in support of one another can yield superior team performance, the presence of even a single disharmonious individual can disrupt group organization and result in substantial performance loss.

To disentangle individual contributions from collective dynamics, we compared fully artificial agent simulations with conditions in which one agent was replaced by a human or an artificial agent with different properties. In cognitive and behavioral sciences, humans are often assumed to flexibly adapt their behavior to context, frequently outperforming simple rules or heuristics^24,25^. Accordingly, one might expect human participants to adjust their strategies depending on environmental conditions and thereby enhance team performance. Unexpectedly, however, we found that teams including a human attacker consistently underperformed relative to teams composed solely of artificial agents implementing simple heuristics. Our results demonstrate that even a single discordant individual, cognitively and motorically constrained–that is, boundedly rational–can destabilize collective coordination and substantially degrade overall team performance.

## Results

Participants, acting as the red attacker, were asked to make as many passes as possible in a dynamic, interactive triangular passing task (Fig. 1a). Each attacking player could choose among three actions, regardless of whether it was in possession of the ball or not. When on-ball, the options were “pass to the left teammate,” “pass to the right teammate,” or “retain possession.” At the moment of passing, the ball was directed toward the current position of the intended receiver. When off-ball, the options were “move left,” “move right,” or “stay in place.” The attacking agents were modeled following simple heuristics for intuitive implementation (Fig. 1b). Specifically, in on-ball cases, the attacking agent compared the left and right pass angles (*θ_l_* and *θ_r_*) and passed to the side with the larger angle, unless the defender was too far away. In off-ball cases, the agent moved to widen its passing angle until the passing lane reached a maximum threshold angle (Max*θ*), unless it was already too close to the ball carrier. In this study, the maximum threshold angle was set to 50 degrees. The defensive agent was modeled using deep reinforcement learning to enable realistic interactions (Fig. 1c). The defensive agent was trained through repeated play against the artificial attacking agents to learn to intercept passes (see Methods for details). The trained defensive agent was pre-tested against human players with experience in various sports and was confirmed to be fairly strong (see Supplementary Videos).

**Figure 1.**
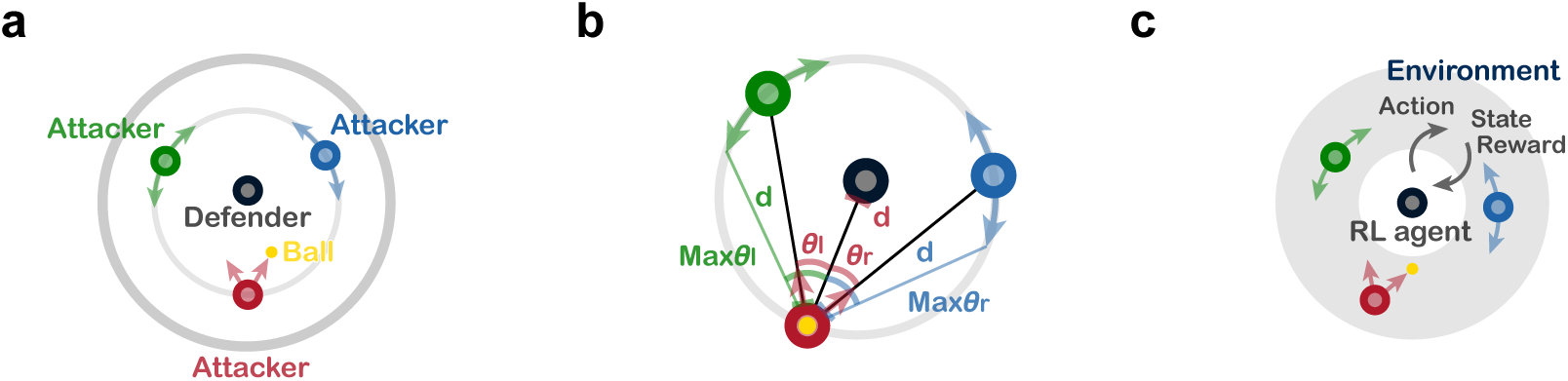
Experimental setup and models. (a) Triangular passing task. Three attackers attempted to maintain possession against a single defender. The ball was passed around the circle, while the defender attempted to intercept. (b) Attacker model. The attacking agents followed heuristic rules such as passing to the wider angle and maintaining viable passing lanes for teammates. (c) Defender model. The defensive agent was trained using deep reinforcement learning to intercept passes.

To manipulate task difficulty, we varied the movement speed of both the defender and the ball at three levels (low, medium, high), resulting in nine experimental conditions in total. The same attacking and defensive agents were used across all conditions. Teams including a human attacker achieved fewer successful passes than teams composed solely of artificial attacking agents (Fig. 2a,b). Scatter plots comparing the two clearly showed that, across all conditions, the data points lay above the unity line, indicating systematic underperformance of humans relative to simple heuristics (Fig. 2c; mixed-effects model, *z* = 13.35, *p* < 0.001).

**Figure 2.**
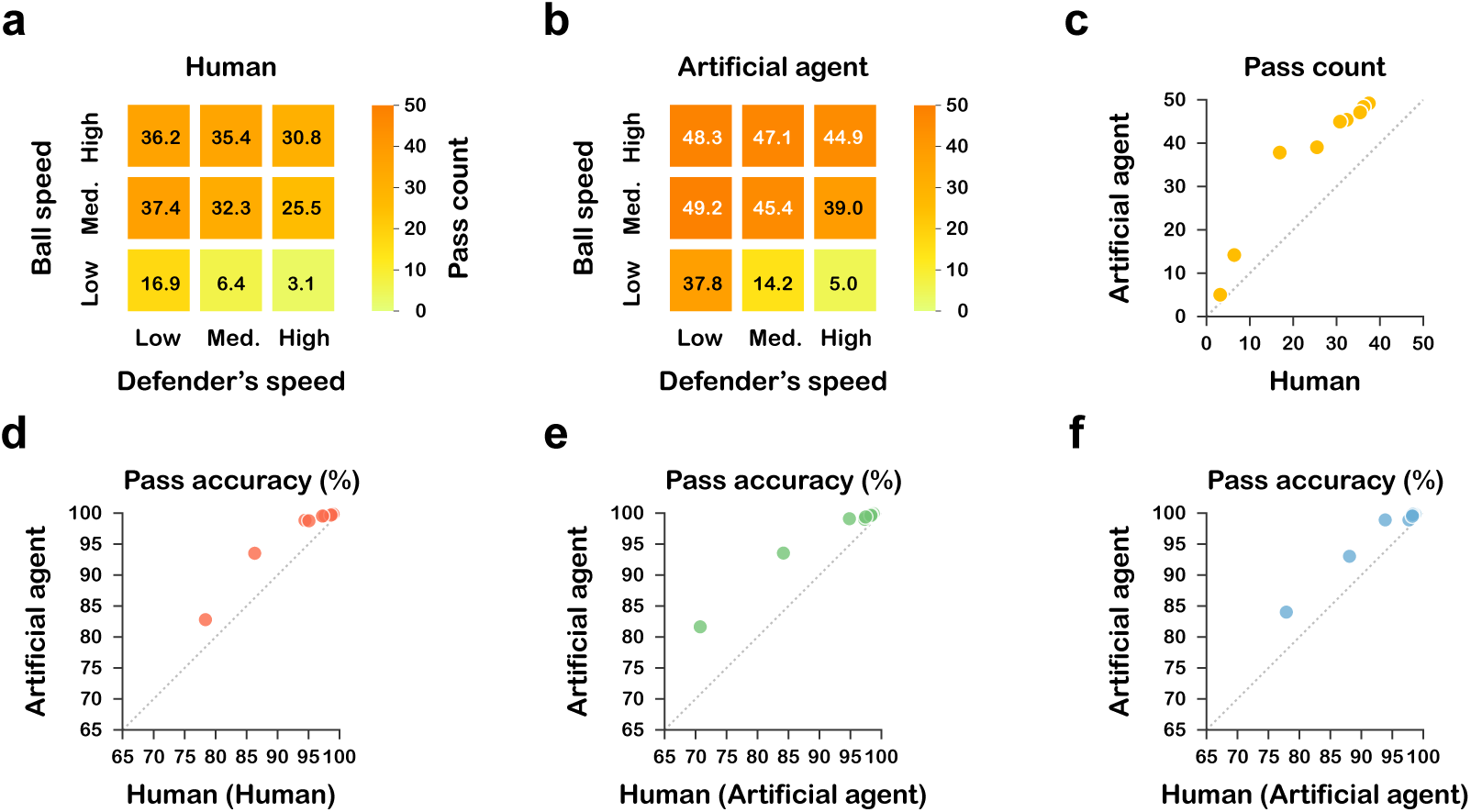
Performance of humans and artificial agents. (a) Average number of successful passes achieved by teams including a human participant across nine experimental conditions (ball speed × defender speed). (b) Average number of successful passes achieved by teams composed entirely of artificial agents under the same conditions. (c) Average number of successful passes between human-including teams and all-agent teams. Each point represents one condition. (d–f) Pass success rates between human-including teams and all-agent teams across conditions. Note that in the human-including teams, the red agent was controlled by a human, whereas the green and blue agents followed the same heuristic rules as the green and blue agents in the all-agent teams.

To investigate the underlying factors of this performance gap, we analyzed the pass success rates of each attacker and found reduced success not only in the human-controlled red attacker but also in the green and blue artificial attackers (Fig. 2d–f). Because the green and blue attackers were governed by identical fixed rules, their performance decline can be attributed to the behavior of the human participant, indicating that individual actions indirectly reduced teammates’ success rates (red: mixed-effects model, *z* = 19.92, *p* < 0.001; green: *z* = 9.30, *p* < 0.001; blue: *z* = 10.01, *p* < 0.001).

To better understand how this occurred, we analyzed off-ball behavior (Fig. 3a). We found that human participants generally exhibited less movement and tended to remain within a restricted area compared with artificial agents (Fig. 3b,c, in contrast to d). To clarify the impact of restricted off-ball movement on performance, we manipulated the maximum pass angle threshold (Max*θ*) in the red artificial agent. Reducing this threshold qualitatively reproduced the constrained off-ball behavior observed in humans (Fig. 3e,f), and progressively reducing the Max*θ* of the red agent revealed that such restrictions propagated across players and degraded teammates’ performance (Fig. 3g; green: mixed-effects model, *z* = 2.15, *p* = 0.031; blue: *z* = 5.25, *p* < 0.001). In contrast, the pass success rate of the red agent itself slightly improved under these conditions (Fig. 3g; red: *z* = −9.04, *p* < 0.001).

**Figure 3.**
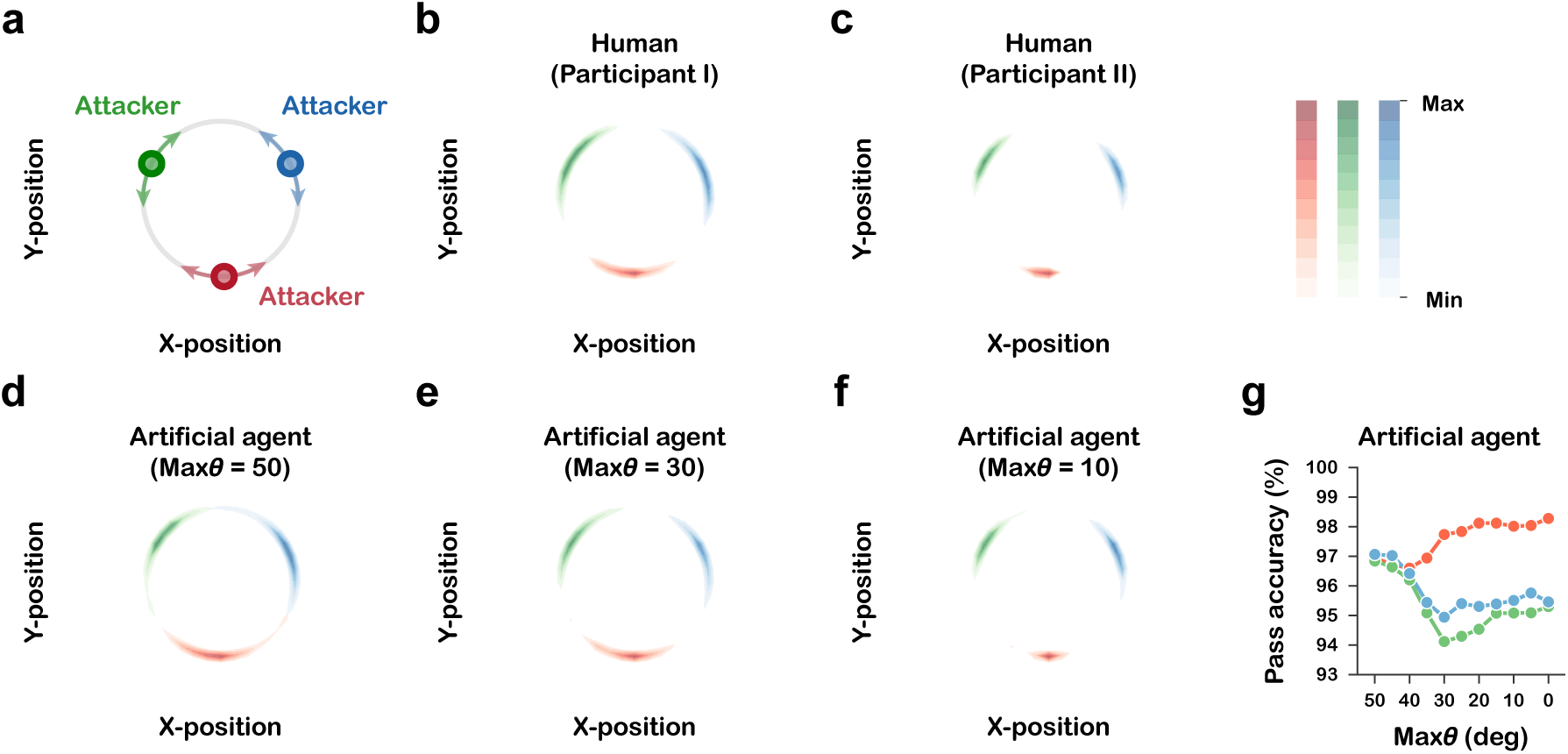
Off-ball movement and its impact on team performance. (a) Schematic illustration of the off-ball movement of the three attacking agents. (b,c) Heatmaps of off-ball movement in human-including teams (two representative participants). (d–f) Heatmaps of off-ball in all-agent teams in which the red agent was assigned different values of maximum pass angle thresholds (Max*θ* = 50, 30, 10). (g) Pass success rates of each agent under different values of Max*θ* assigned to the red agent in the all-agent teams. Each line represents one of the three agents (red, green, or blue).

Next, to further investigate how teammates’ performance was degraded, we examined the dynamical patterns formed by the attackers. Specifically, we focused on the internal angles of the triangle defined by the positions of three attacking players (Fig. 4a) and visualized their dynamics by plotting trajectories in a phase plane. As expected, when a human participant with a restricted movement range was included in the team, the dynamic patterns exhibited reduced symmetry compared with those of teams composed solely of artificial agents (Fig. 4b,c, in contrast to d). Using the time series of the internal angles of the triangle, we quantified the occurrence of three dynamic coordination patterns—“Rotation,” “Partial anti-phase,” and “Partial in-phase” (Fig. 4g–i; see Methods for details). We found that as Max*θ* decreased, the frequency of the Rotation pattern declined (Fig. 4j; Rotation: mixed-effects model, *z* = 23.9, *p* < 0.001), whereas the frequency of the Partial anti-phase pattern increased (Fig. 4j; Partial anti-phase: *z* = −32.6, *p* < 0.001; Partial in-phase: *z* = −0.07, *p* = 0.94). Previous studies of football players have reported a similar ordering of coordination patterns: in skilled groups, the dominant pattern is Rotation, followed by Partial anti-phase and then Partial in-phase, whereas in less skilled groups, the order of the first two patterns is reversed, with Partial anti-phase becoming most frequent^20^. Consistent with this finding, our results indicate that when Max*θ* was around 50 degrees, teams exhibited well-organized cooperative patterns dominated by Rotation. However, when Max*θ* fell below approximately 40 degrees, the coordination structure shifted toward less organized patterns, characterized by more Partial anti-phase occurrences. This demonstrates that the inclusion of even a single disharmonious agent can disrupt collective order and degrade the overall quality of team coordination.

**Figure 4.**
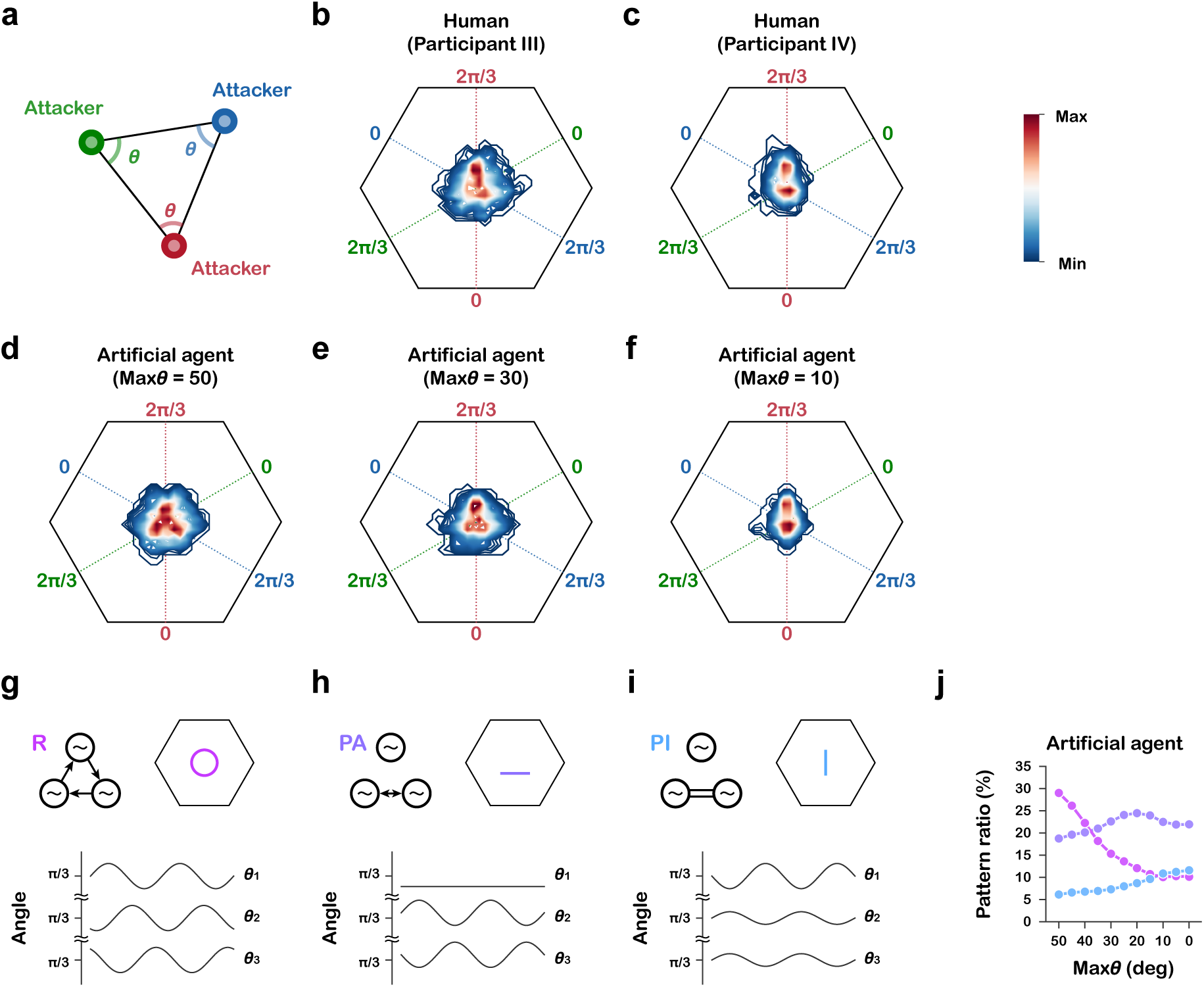
Dynamic coordination patterns of attackers. (a) Schematic illustration of the internal angles of the triangle defined by the three attackers (*θ*_1_, *θ*_2_, *θ*_3_). (b,c) Phase-plane heatmaps of internal angles in human-including teams (two representative participants). (d–f) Phase-plane heatmaps of internal angles in all-agent teams in which the red agent was assigned different values of maximum pass angle thresholds (Max*θ* = 50, 30, 10). (g–i) Examples of three dynamic coordination patterns: Rotation (R), Partial anti-phase (PA), and Partial in-phase (PI), illustrated with schematic diagrams and corresponding time series of internal angles. (j) Pattern ratio of each coordination pattern under different values of Max*θ* assigned to the red agent in all-agent teams. Each line represents one of the three patterns (magenta, purple, or cyan).

Since constraints on off-ball movement alone could not account for the reduction in human participants’ own pass success rate, we next focused on decision-making when in possession of the ball. Specifically, we focused on pass angles and actual passing choices (Fig. 5a) and found that human passing did not exhibit the sharp separation between options that is characteristic of deterministic heuristic agents; instead, participants frequently chose to pass to a teammate with a narrower angle (Fig. 5b,c in contrast to h). Although various cognitive and strategic factors may underlie this tendency, we investigated participants’ decision process with reference to the framework of diffusion decision models, which describe human decision-making as evidence accumulation under noise^26–28^. We first calculated the probability of passing to the green agent for each Δ*θ* (*θ_l_ − θ_r_*) and found that, overall, human participants tended to pass to the teammate with the wider pass angle (Fig. 5d). However, unlike typical two-alternative decision tasks^29,30^, there was no increase in response time in more difficult situations (i.e., when the absolute Δ*θ* was small). Instead, response times remained constant at approximately 250 ms across Δ*θ* (Fig. 5e), indicating that participants typically selected their passing option with little deliberation and passed immediately after receiving the ball (see also Fig. S1). Furthermore, there was little left–right bias in passing frequency (Fig. 5f). Given that human participants intended to pass to the teammate with the wider angle, the sharpness of separation in their choices could provide insight into the sensory information they relied on. For instance, if participants first selected the passing direction and then executed the motor action in a sequential manner, a sensorimotor delay of several hundred milliseconds would be expected^31^. In such a case, using information from slightly earlier than the actual pass might better account for their behavior, yielding a clearer separation. To test this, we recalculated passing probabilities using Δ*θ* taken at earlier time points (with one time-step corresponding to 100 ms) relative to the pass event at time *t* (Fig. 5g). Our results showed that separation became less distinct when older information was used, suggesting that decisions were made based on the most recent information available before the pass. Additionally, we examined models in which sensory evidence was accumulated over time, including leaky accumulation that placed greater weight on recent information. Across these variants, however, sharper separation in passing choices was observed when decisions relied primarily on the most recent information available before the pass (Fig. S2). Taken together, our results consistently suggest that participants based their passing decisions on sensory information immediately preceding the pass. Since time *t* refers to the moment of passing, and considering delays in the human sensorimotor system, their decisions were likely based on predicted future states. Building on these analyses, we modeled the decision process as relying on information immediately preceding the pass, subject to noise. Here, “noise” can be interpreted broadly, encompassing both internal noise in the sensorimotor process and errors in predicting pass angles. The red agents whose passing decisions were made following summation of Δ*θ* and zero-mean Gaussian noise with variance *σ*^2^ qualitatively reproduced the passing choice tendencies observed in humans when *σ* was increased (Fig. 5i,j). Moreover, increasing *σ* decreased the pass success rate of the red agent itself (Fig. 5k; red: mixed-effects model, *z* = −4.74, *p* < 0.001), while having little effect on the pass success rates of its teammates (Fig. 5k; green: *z* = −0.28, *p* = 0.78; blue: *z* = 0.39, *p* = 0.70).

**Figure 5.**
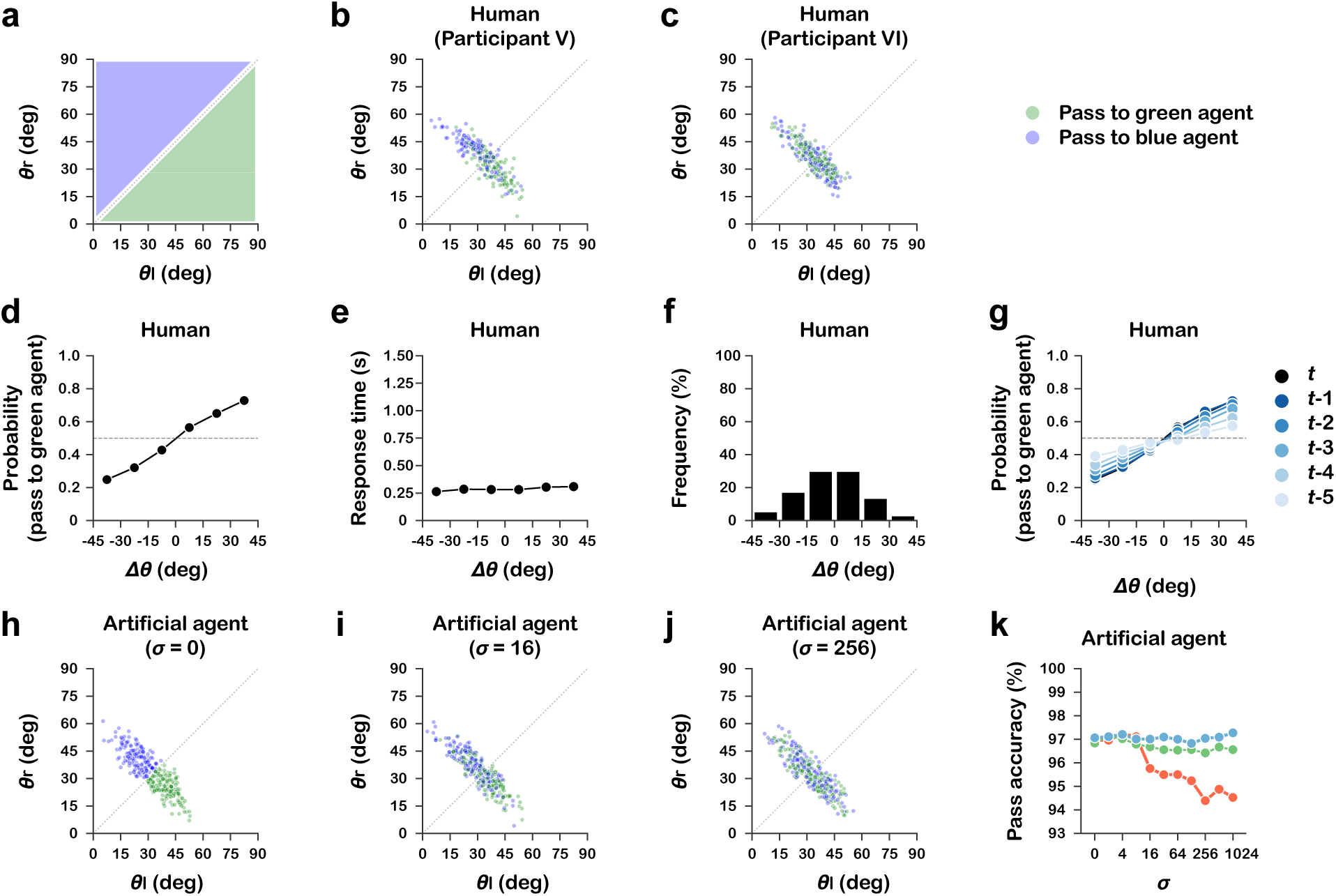
Decision-making when in possession of the ball. (a) Schematic illustration of passing options. The green region indicates where the left passing angle (*θ_l_*) is larger than the right passing angle (*θ_r_*), and the blue region indicates where *θ_r_* is larger than *θ_l_*. (b,c) Passing choices in human-including teams (two representative participants). (d) Probability of a human passing to the green (left-side) agent for each Δ*θ* (*θ_l_*– *θ_r_*). (e) Response time from receiving a pass to making the next pass for each Δ*θ*. (f) Relative frequency of passes for each Δ*θ*. (g) Probability of a human passing to the green agent for each Δ*θ*, computed using information from different time points. (h–j) Passing choices of in all-agent teams in which the red agent was assigned different levels of noise (*σ* = 0, 16, 256). (k) Pass success rates of each agent under different levels of *σ* assigned to the red agent in the all-agent teams. Each line represents one of the three agents (red, green, or blue).

Finally, we examined the extent to which the performance of groups including a human participant could be reproduced by integrating the results obtained thus far. Specifically, for each human participant, we estimated Max*θ* and *σ* from their off-ball and on-ball behaviors (see Methods and Fig. S3 for details), and then compared the performance of simulations using the estimated values with the performance of groups including a human participant. We found that the performance of human-including groups was reproduced more accurately across the nine experimental conditions, although the simulated performance still lay slightly above the unity line (Fig. 6a; mixed-effects model, *z* = 13.35, *p* < 0.001). Moreover, reproduction was not limited to overall team performance; the pass success rates of the individual red, green, and blue agents were also accurately reproduced (Fig. 6b–d; red: *z* = 2.21, *p* = 0.027; green: *z* = 3.31, *p* < 0.001; blue: *z* = 7.32, *p* < 0.001). These findings provide evidence that the off-ball and on-ball parameters we focused on captured key characteristics of human behavior.

**Figure 6.**
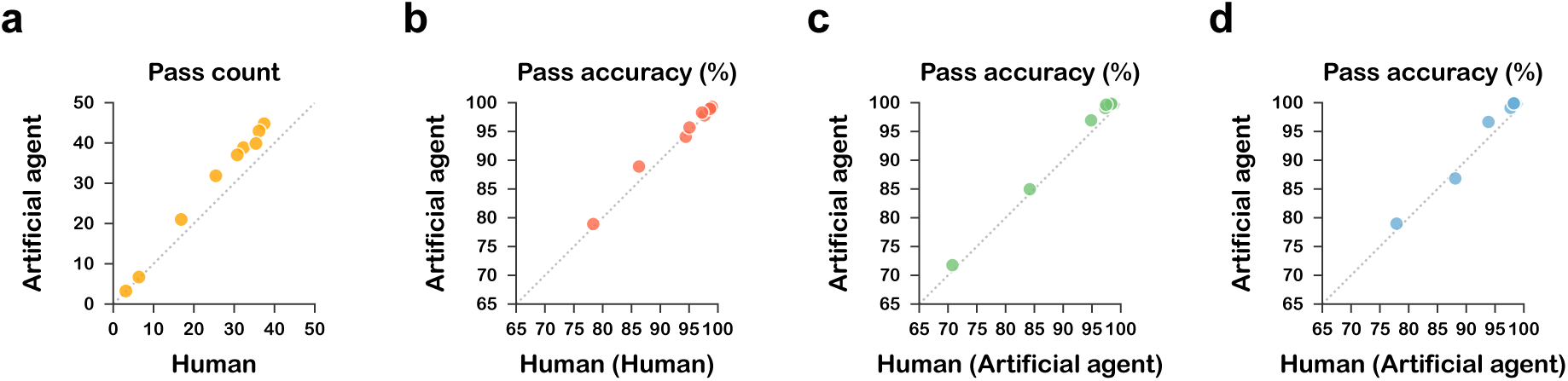
Reproducing human performance with integrative modeling. (a) Average number of successful passes between human-including teams and all-agent teams in which the red agent is parameterized by human-derived estimates. Each point represents one condition. (b–d) Pass success rates between human-including teams and all-agent teams in which the red agent is parameterized by human-derived estimates.

## Discussion

A long-standing view in cognitive science and behavioral research is that humans adaptively select actions that enhance team performance, often performing as well as or better than simple rules or heuristics^24,25^. Our findings, by contrast, show that, in the dynamic, cooperative setting of the triangular passing task, humans fell short of artificial agents implementing simple heuristics. Notably, the presence of a human participant not only reduced their own success rate while in possession but also indirectly lowered that of their teammates, likely due to the propagation of suboptimal choices. These results suggest that individual decisions are not always adaptively tuned, and that such suboptimality can cascade across teammates, ultimately degrading cooperative performance.

One of the key contributions of this study is that it quantitatively demonstrates how individual constraints can propagate to others and thereby undermine collective performance. Across diverse fields such as cognitive, social, and sports sciences, it has long been recognized that the effectiveness of groups depends not only on the apparent contributions of individuals but also on more subtle, often hidden behaviors that shape coordination^32–34^. Yet, in natural settings, it is generally difficult to disentangle and measure the extent to which the actions of a single individual influence overall group outcomes. By introducing a cooperative task involving humans and artificial agents and comparing the outcomes with counterfactual simulations, this study provides clear evidence that individual limitations can trigger cascades that compromise collective performance within the dynamics of cooperation.

Our results are consistent with symmetry-based theoretical frameworks, which predict that collective coordination converges to a limited set of dynamic patterns^35,36^. Approaches based on geometrical symmetries, originally developed to explain intra-individual coordination such as locomotor gait patterns^37–40^, have more recently been extended to interpersonal coordination^20^. In this context, previous work with football players has demonstrated that skilled groups predominantly exhibit the Rotation pattern, whereas less-skilled groups more often display the Partial anti-phase pattern, suggesting that skill level functions as a bifurcation parameter in human synchronization^20^. Our findings extend this view by showing that when even a single individual’s movement was constrained, collective dynamics shifted from the more symmetric Rotation pattern toward less symmetric Partial coordination patterns.

In this study, we were able to more accurately reproduce human-like behavior and team performance by introducing artificial agents based on human tendencies in both off-ball and on-ball actions. Although the simulation using the estimated parameters provided a close match to the human data, these findings should be interpreted with caution. The passing choices of participants, which were explained here in terms of decision-making under noise, may also have contained elements of strategic intent. For example, skilled players might sometimes take the risk of attempting a narrower passing lane if they judged it could create a more favorable situation for the team. Given that the performance of human-including teams did not surpass that of deterministic agents, the likelihood that participants employed such sophisticated cognitive strategies is relatively low. Nevertheless, further investigation will be required–especially with regard to human decision-making while in possession and potential behavioral adaptations through learning^41,42^–to achieve a more comprehensive understanding.

In conclusion, our results demonstrate that limitations of individual rationality can propagate through teammates, destabilizing collective coordination and reducing group performance. Moreover, by incorporating these constraints into artificial agents, we were able to more accurately reproduce human-like behavior as well as the decline in performance observed when humans were included. These findings suggest that disruption introduced by even a single individual can dramatically alter, and at times destabilize, collective dynamics. By highlighting the link between individual and collective dynamics across both humans and artificial agents, this study provides useful insights not only for understanding coordination in social and organizational contexts, but also for developing adaptive human–AI systems^43,44^.

## Methods

### Participants

Twenty-eight participants took part in the experiment (2 females, 3 left-handed, aged 18–24, mean = 20.79, s.d. = 1.47). Participants varied in their prior experience with sports: some had extensive experience in activities such as football or basketball, whereas others had little experience beyond physical education classes (Table S1). This study was approved by the Ethics Committee of Ritsumeikan University (BKC-LSMH-2021-088-3) and Nagoya University (hc21-08). Informed consent was obtained from each participant before the experiment. Participants received JPY 1,000 per hour as a reward. To promote sustained task engagement, deceptive instructions indicated that an extra JPY 2,000 would be paid if they achieved a higher number of successful passes, without disclosing any specific numerical criterion. Participants were fully debriefed after the experiment and received the additional JPY 2,000, independent of their task performance.

### Task design

We used a cooperative triangular passing task, inspired by team ball sports^20–23^. All positions were represented in PsychoPy normalized screen coordinates, where the two-dimensional plane was centered at the origin and spanned from −1 to 1 along both the *x*- and *y*-axes. The playing field was depicted as a circular boundary (dark gray in Fig. 1a) with a radius of 0.8. Attackers were constrained to move along an inner circle of radius 0.5 (light gray in Fig. 1a), whereas the defender could move freely inside the field. Each player and the ball were displayed as circular disks: the diameter was 0.10 for the attackers and defender, and 0.05 for the ball. At the beginning of each episode, the three attackers (red, green, and blue) were placed at the vertices of an equilateral triangle on the circle (e.g. red at -*π*/2, green at 5*π*/6, blue at *π*/6), and the ball was randomly assigned to one of them. The defender started from a random position within the range of −0.3 to 0.3 along both the *x*- and *y*-axes. The ball carrier (red agent in Fig. 1a) had three on-ball actions: to pass to the left teammate (green agent), to pass to the right teammate (blue agent), or to retain possession. For simplicity, a pass was always directed toward the instantaneous position of the intended receiver. Off the ball, attackers had three actions: move left (clockwise) along the circle, move right (counterclockwise), or stay in place, thereby attempting to secure passing lanes for the teammate with the ball (see Heuristic agents for details). The defender moved to intercept the ball. A catch or interception was registered when the Euclidean distance between an agent and the ball fell below 0.10, which approximately corresponded to visual “contact” on the display. An out-of-bounds event occurred when the ball traveled beyond a radial distance of 0.8, that is, when a pass was not caught by an agent and the ball contacted the outer circular boundary. Each episode terminated when one of the following conditions was met: (i) the attackers successfully completed 50 passes, (ii) the defender intercepted a pass, or (iii) the ball went out of bounds.

### Experimental conditions

Task difficulty was manipulated by varying ball speed and defender speed at three levels (low, medium, high), yielding nine conditions in total. The defender followed a pretrained policy (see RL agent subsection), and the levels were determined through pre-experiments so that task performance spanned a wide range—from easy (passes almost always succeeded) to difficult (passes often failed). Each participant performed eight trials per condition, repeated across three sets, resulting in 216 trials in total. The order of conditions was counterbalanced across participants. To become familiar with the task, participants first completed three practice sets under the medium–medium condition. The position of the defensive agent was computed by integrating the acceleration (i.e., the selected action) twice with the Euler method, with viscous resistance proportional to velocity applied to moving agents^45,46^. Attacking agents followed the same scheme but were projected back onto the circle of radius 0.5 at each step to remain constrained to the ring. By contrast, the ball moved with piecewise constant velocity: when passed, its velocity was set along the pass direction and then maintained without viscous drag until reception or interception. The low, medium, and high conditions corresponded to the following movement scales (arbitrary unit, a.u.): ball = 3.6, 4.8, 6.0; defender = 1.5, 1.75, 2.0; attackers = fixed at 2.0. Because the mass of the ball (0.3) was smaller than that of the agents (1.0), the ball could travel substantially faster than the players.

### Heuristic agents

The attacking artificial agents acted according to deterministic heuristics (Fig. 1b). Each attacking agent had three discrete actions available both on-ball and off-ball. When on-ball, the options were “pass to the left teammate,” “pass to the right teammate,” or “retain possession.” When off-ball, the options were “move left,” “move right,” or “stay in place.” The decision rules for attacking agents were as follows:

### On-ball: Passing rules

1. If the distance between the ball holder and the defender exceeded a preset threshold (2.0 a.u.), the ball holder did not pass and retained possession (red *d* in Fig. 1b).
2. If the distance was below the threshold, the two passing angles (*θ_l_* and *θ_r_* in Fig. 1b) were evaluated, and the ball was passed to the teammate with the larger angle.
3. However, to avoid reckless passes, if both passing angles were smaller than a minimum threshold (30^◦^), the ball was retained.

### Off-ball: Positioning rules

1. Off-ball players moved left or right along the circle (light gray line in Fig. 1b) to widen the passer’s angle (*θ*) until the passing angle reached the maximum threshold (Max*θ* in Fig. 1b).
2. To preserve the triangular formation, if the distance to the ball holder or the other potential receiver became too small (below 0.85 a.u.), the agent remained in place (green and blue *d* in Fig. 1b).
3. If a pass was directed toward them, the player moved to the intersection point of the pass trajectory and the circular path, waiting to receive the ball.

These constitute the basic behavioral rules. The same set of rules was applied consistently across all nine experimental conditions. However, to better reflect the characteristics of human behavior, the rules for the red agent were later modified, with a restricted movement range when off-ball and a probabilistic choice when on-ball, as detailed in the subsection on Computational simulation.

### RL agent

The defender was modeled as a deep reinforcement learning (RL) agent trained to intercept passes. We adopted a value-based approach formalized as a Markov decision process (MDP)^47^, where the agent learns to maximize expected discounted returns by estimating an action-value function *Q*(*s*, *a*). Formally, the optimal action-value function is given by the Bellman equation:

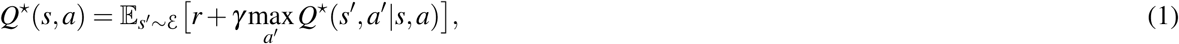

where *s*^′^ and *a*^′^ denote the next state and action, respectively, and *γ* is the discount factor. To approximate *Q*(*s*, *a*), we used a deep Q-network (DQN)^48^ and incorporated several extensions that address well-known limitations of DQN: double DQN to reduce overestimation bias^49^, prioritized experience replay to improve sample efficiency^50^, and a dueling network architecture to stabilize value estimation^51^. The training objective was to minimize the temporal-difference error, yielding the following loss function:

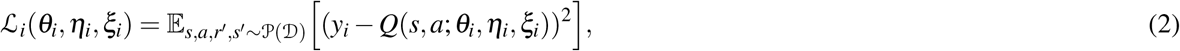

Where

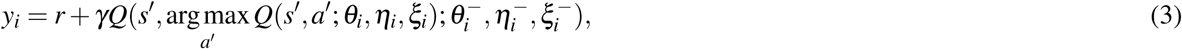

and *P*(·) denotes prioritized sampling from the replay buffer. Here, *θ* represents the parameters of the common layers, while *η* and *ξ* are the parameters of the value and advantage streams in the dueling network, respectively. The parameters *θ_i_*^−^, *η_i_*^−^, *ξ_i_*^−^ denote those of the corresponding target network. The input to the Q-network consisted of the positions and velocities of the ball, the defender itself, and the three attackers, represented in both absolute and relative coordinates, informed by previous work in ethology and neuroscience^52–54^. We assumed that sensory delays were compensated for by internal estimation of self- and other-motion^55–57^, and thus the current information at each time step was used as input. The output layer corresponded to discrete movement actions in 12 directions, separated by 30^◦^, aligned with the design used in previous studies of multi-agent interactions^58,59^. The Q-network was a multilayer perceptron with two hidden layers of 64 units each, followed by two separate streams of 32 units implementing the dueling architecture^51^. ReLU activations^60^ were used throughout. Network parameters were optimized using the Adam optimizer^61^ with Huber loss^62^. The replay memory size was 10^4^, minibatch size was 32, and the learning rate was 10^−6^. The discount factor *γ* was set to 0.9, and the prioritized replay exponent *α* was 0.6. An *ε*-greedy exploration policy was employed, with *ε* linearly annealed from 1.0 to 0.1 over the first 10^4^ episodes and fixed at 0.1 thereafter. Training was conducted for 3 10^6^ episodes under the following environment settings. The movement scales were set to 3.0 for the ball, 2.0 for the defender, and 2.0 for the attackers. For the attacking agents, the minimum distance to the defender for passing was set to 1.0, so that the ball holder retained possession when the defender was sufficiently far away. The minimum passing angle was set to 30°, so that the ball holder retained possession until a receiver created a passing lane. To generate more diverse situations, zero-mean Gaussian noise with a variance of 0.05 was added to each pass. The maximum movement angle was set to 45°, so that off-ball players moved along the circular path to widen the passer’s angle until the passing angle reached this threshold. The minimum distance to the passer was set to 0.8, so that off-ball players moved away from the passer to avoid getting too close. For the defensive agent, a reward of +1 was given for successfully intercepting the ball (i.e., when the distance to the ball became less than 0.1), and a penalty of –1 was given when it moved out of bounds. Each episode terminated either when a reward was obtained or when 300 time steps had elapsed, after which a new episode was initiated. Target-network parameters were updated every 2000 episodes. In the human and simulation experiments, *ε* was set to 0 and the RL agent acted greedily according to the learned policy.

### Human Experiment

In the human experiment, only the red agent among the attacking agents was controlled by human participants. Participants were seated in a chair and instructed to “make as many passes as possible.” They controlled the red agent on the screen using the two central buttons of an Xbox One controller. When in possession of the ball, pressing the left button passed to the left teammate (green agent), pressing the right button passed to the right teammate (blue agent), and pressing neither button kept possession. When not in possession, pressing the left button moved the red agent clockwise along the circle, pressing the right button moved it counterclockwise, and pressing neither button kept the agent in place. To allow for more intuitive control, the initial positions of the agents were fixed such that the red agent started at the bottom of the screen ( − *π*/2), the green agent at the upper left (5*π*/6), and the blue agent at the upper right (*π*/6). The stimuli were presented on a 15.6-inch monitor at a refresh rate of 240 Hz. To ensure the stable real-time presentation of defensive agents controlled by deep reinforcement learning, the position and velocity of each agent on the screen were updated at 10 Hz, and positions during each episode were recorded at 10 Hz on a computer (GALLERIA UL7C-R36, Thirdwave Corp, Japan) using PsychoPy version 2022.1.3. The viewing distance of the participants was approximately 60 cm.

### Computational simulation

For each condition, we ran 28 independent simulations, matching the number of human participants, by varying random seeds (i.e., the initial position of the defender). In the first set of simulation experiments, we used the heuristic agents described above, with the red agent set to the same parameters as the green and blue agents (Max*θ* = 50, *σ* = 0; Fig. 2). Next, to introduce a restriction on the movement range of the red agent, Max*θ* was decreased from 50 to 0 in steps of 5 degrees, resulting in 11 simulation levels (with *σ* fixed at 0; Fig. 3). Subsequently, we introduced a noisy decision process for the red agent. We assumed that the passing decision was based only on the information available at the instant of passing, rather than on accumulated evidence across multiple frames. Specifically, at time *t* the noisy passing evidence was defined as

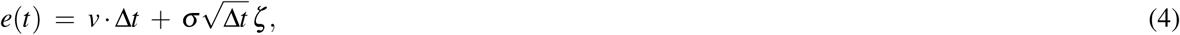

where *v* denotes Δ*θ* (*t*), the left–right passing angle difference at time *t*, Δ*t* is the frame interval (100 ms), *σ* is the noise scale, and *ζ* ∼ *N*(0, 1) is a Gaussian random variable. The decision rule was then simply

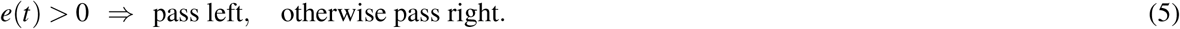

Here, *σ* was varied from 0 to 1024, resulting in 11 simulation levels (with Max*θ* fixed at 50; Fig. 4). The starting point was fixed at 0, corresponding to no initial bias. Thus, this is essentially equivalent to probabilistic passing based on the Boltzmann choice rule defined by the following equation^47^.

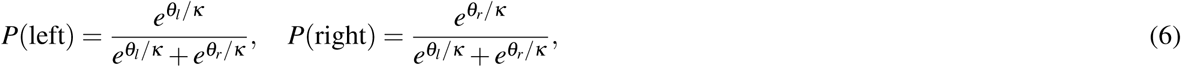

where *θ_l_* and *θ_r_* denote the left and right passing angles, respectively, and *κ* is a parameter controlling the degree of uncertainty. Intuitively, as *κ* → 0 the choice becomes nearly deterministic, always favoring the larger angle, whereas as *κ* → ∞ the choice approaches a random 50–50 distribution (Fig. S4).

### Model fitting

We conducted model fitting designed to incorporate the behavioral characteristics of human participants. A participant’s Max*θ* was estimated from comparisons with the red agent’s angular position distributions across different Max*θ* values. Specifically, all positions were converted to polar coordinates and binned into 36 angular bins along the circle. From the resulting histogram we obtained a probability vector 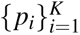 and computed the normalized entropy

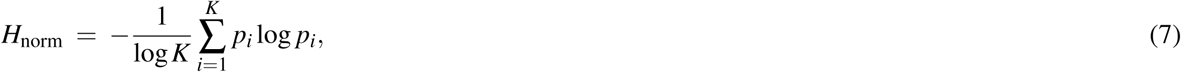

with *K* = 36 and natural logarithms. Intuitively, *H*_norm_ = 0 if the agent never moves (all positions fall into a single bin), and it increases as movement becomes more variable, reaching *H*_norm_ = 1 when the distribution is perfectly uniform. We computed *H*_norm_ for each simulated Max*θ* and for each participant, and estimated a participant’s Max*θ* by minimizing the mean absolute error (MAE) between simulation and human values. Because Max*θ* was simulated in increments of 5°, values between the discrete levels were linearly interpolated before fitting. A participant’s *σ* was estimated from the separation index of passing decisions. Let *θ_max_* be the larger of the two available passing angles at the decision moment and *θ_chosen_* be the angle corresponding to the actually chosen pass. The separation index was defined as

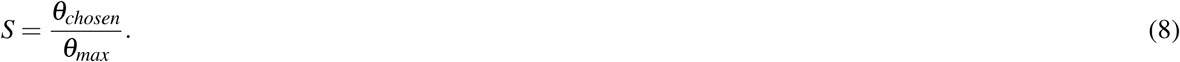

Thus, *S* = 1 indicates that the wider angle was always chosen, whereas *S* ≈ 0.5 indicates approximately random choice between the two receivers. We computed *S* for each simulated *σ* and for each participant, and estimated a participant’s *σ* by minimizing the MAE between simulation and human values. Because the simulated raw separation index values from the simulated data were somewhat noisy, linear interpolation was not applicable. Instead, we fitted the data with a monotonically decreasing exponential function, which was then used to estimate the participant’s *σ* values. Notably, Max*θ* and *σ* were estimated from distinct behavioral features, and their effects were found to be nearly independent (Fig. S3a,b). Most participants released the ball immediately after receiving it; however, in practice there was typically a delay of about 1–3 frames, which was likely due to factors related to controller operation (Fig. S1). In other words, the delay could serve to prevent the agent from moving prematurely and failing to catch the ball. To capture this tendency, we modeled the probability of releasing a pass using a geometric distribution with a mean of 2.5 (i.e., pass probability 0.4). For simplicity, the delay parameters were constrained to be identical across all participants and conditions. The final reproduction of human data was obtained with parameters Max*θ* = 20 and *σ* = 32, which were closest to the mean of the estimated participant values (Fig. S3c).

### Data analysis

All analyses were conducted in Python 3.8. The pass count was defined as the total number of successful passes completed before the episode ended (maximum 50 per episode). Pass accuracy was defined as the proportion of successful passes out of all passes attempted. To capture the group-level dynamics, we analyzed the temporal evolution of the triangle formed by the three attackers. At each frame, the interior angles (*θ*_1_, *θ*_2_, *θ*_3_) were computed from their positions. These time series were converted into phase variables by detecting successive extrema and linearly interpolating the phase between them, following previous work on coupled oscillator systems^20^. The *φ_i_*(*t*) denote the phase of attacker *i* (*i* = 1, 2, 3). The phase differences between all pairs were then computed as

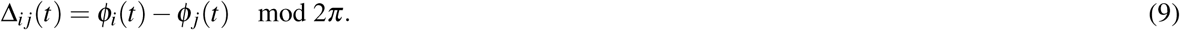

Symmetric Hopf bifurcation theory predicts five generic synchronization patterns for a ring of three coupled oscillators: rotation (R), partial anti-phase (PA), partial in-phase (PI), PA’, and PI’. Cases in which no stable relation was observed were classified as other (O). In the R pattern, all three attackers were approximately shifted by 2*π*/3. In the PA pattern, two attackers moved in anti-phase while the third remained nearly constant (amplitude death). In the PI pattern, two attackers moved in-phase while the third was in anti-phase. PA’ and PI’ denote less symmetric variants in which two attackers were anti-phase or in-phase, respectively, while the third behaved independently. Patterns were detected sequentially through each trial, and consecutive segments of the same type were concatenated.

The relative frequency of each pattern was computed for each condition. For each human participant, the probability of passing to the green agent as a function of Δ*θ* was computed as the number of passes directed to the green agent divided by the total number of passes in each bin. The Δ*θ* values were binned from –45° to 45° in 15° intervals. Reaction time was defined as the number of frames between receiving the ball and releasing the pass. Relative frequency was defined as the proportion of passes in each bin relative to the total number of passes. Notably, 98.15% of all passes (45,720 out of 46,584) fell within the –45° to 45° range. To examine the temporal information underlying passing decisions, we recalculated passing probabilities using Δ*θ* taken at different time points relative to the pass event (*t*). One time step corresponded to 100 ms, and values from earlier frames (*t − k*) were substituted for Δ*θ* (*t*). We next considered an accumulation of evidence from *t − k* up to *t*, implemented as the average of Δ*θ* values across the window:

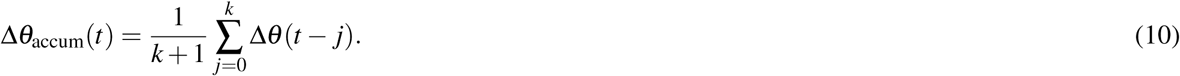

In addition, a leaky accumulation scheme was tested in which recent information was weighted more strongly according to an exponential kernel with time constant *τ*:

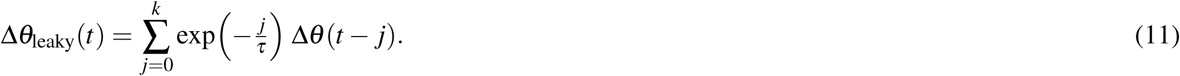

For simplicity, the accumulation window was fixed at five time steps, approximately corresponding to the average time between a teammate releasing the ball and the participant receiving it (Fig. S5).

### Statistical analysis

All statistical analyses were conducted in Python 3.8 using the Statsmodels package. To compare human and agent performance across conditions, we fitted linear mixed-effects models (LMMs) with the mean number of successful passes as the dependent variable. Fixed effects included group (Human vs. Agent), ball speed (three levels), defender speed (three levels), and their interactions. Subjects were treated as random effects with random intercepts and random slopes for ball speed and defender speed. Significance of contrasts (e.g., Human vs. Agent averaged across conditions) was assessed using Wald tests on estimated marginal means (Fig. 2c). For the analyses of pass success rates (Fig. 2d–f), probabilities were logit-transformed prior to model fitting. Separate models were fit for the red, green, and blue attackers, each including the same fixed and random effects structure as described above. For the Max*θ* manipulation analysis (Fig. 3g), logit-transformed pass success rates were modeled as a function of the maximum pass angle threshold, ball speed, and defender speed, with random intercepts and random slopes for threshold, ball, and defender speed across subjects. For the analyses of dynamic coordination patterns (Fig. 4j), the occurrence frequencies of the Rotation, Partial anti-phase, and Partial in-phase patterns were logit-transformed and modeled using LMMs with threshold, ball speed, and defender speed as fixed effects and with random intercepts and random slopes for threshold, ball, and defender speed across subjects. For the noise manipulation analysis (Fig. 5g), pass success rates were logit-transformed and modeled as a function of the noise parameter *σ* (entered on a log_2_-scale to account for its exponential spacing), ball speed, and defender speed, with random intercepts and random slopes for *σ*, ball, and defender speed across subjects. For the reproduction analysis (Fig. 6a–d), mean pass counts and pass success rates of the red, green, and blue agents were analyzed with LMMs including group, ball speed, and defender speed as fixed effects and with random intercepts and random slopes for ball and defender speed across subjects. Models were estimated using restricted maximum likelihood (REML) with the L-BFGS optimizer. The assumption of normally distributed residuals was checked using Quantile–Quantile (Q–Q) plots, which indicated no major deviations from normality. All statistical tests were two-sided, and the significance level was set at *α* = 0.05.

## Data availability

Datasets and the trained model used in this study are available at https://doi.org/10.6084/m9.figshare.30648164.

## Code availability

Code for simulations and figure generation is available at https://github.com/TsutsuiKazushi/pass-game.

## Acknowledgements

This work was supported by the Japan Society for the Promotion of Science (Grants-in-Aid for Scientific Research 22K17673, 24H01434, and 25K02329), the Japan Science and Technology Agency (Strategic Basic Research Programs ACT-X JPMJAX24LA), and the Program for Promoting the Enhancement of Research Universities.

## Author contributions

K.T., N.M. and K.F. conceived the study. K.T. developed and implemented the software. K.T., N.M. and K.F. designed the experiments. N.M. and Y.G. conducted the experiments. K.T. analyzed the data. K.T., N.M., Y.G., R.O., F.K., K.K., T.I., and K.F. wrote the manuscript.

## Supplementary Information

**Figure S1.**
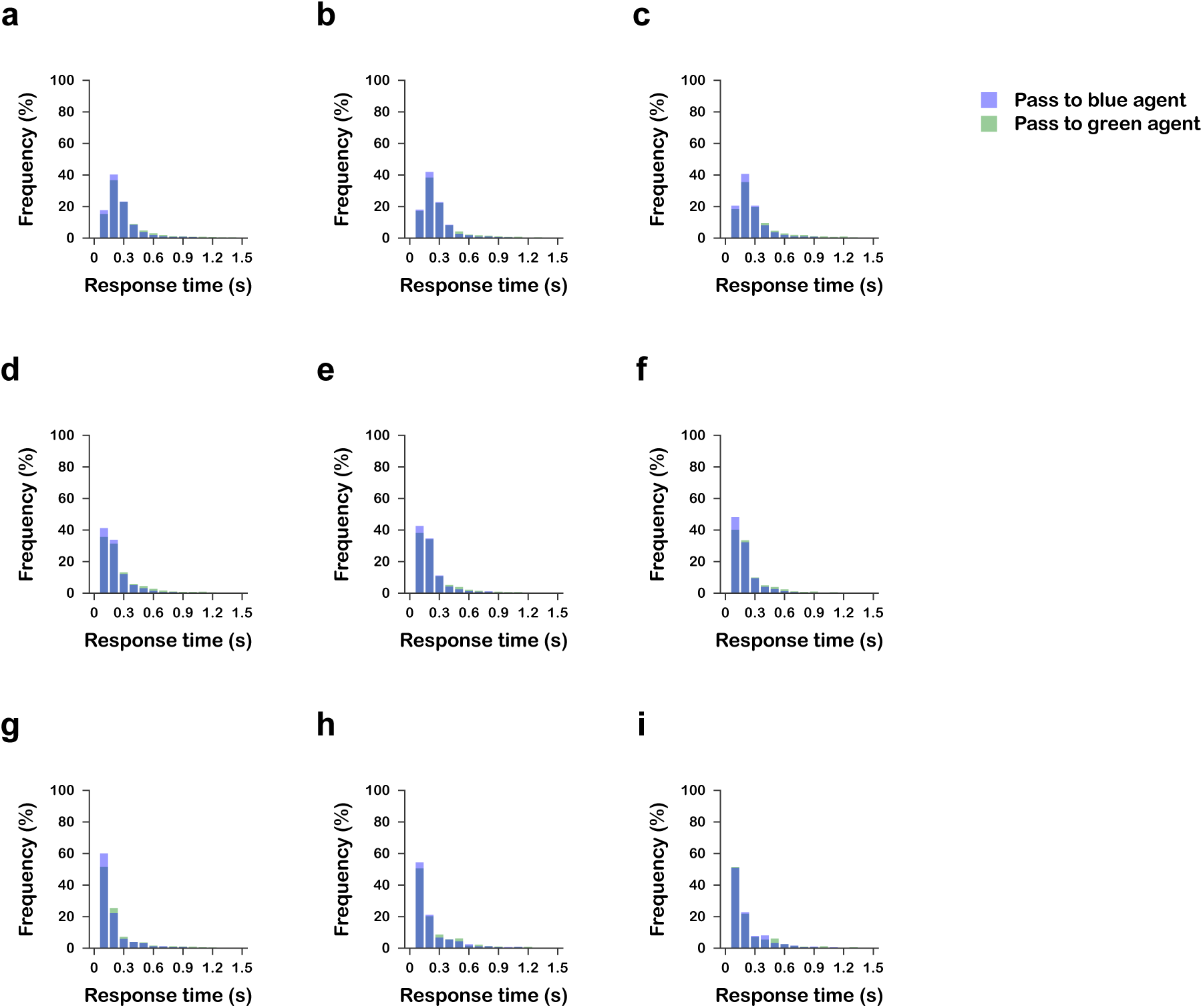
Histograms of response times for passing to each agent across nine experimental conditions. (a) High ball speed, Low defender speed. (b) High ball speed, Medium defender speed. (c) High ball speed, High defender speed. (d) Medium ball speed, Low defender speed. (e) Medium ball speed, Medium defender speed. (f) Medium ball speed, High defender speed. (g) Low ball speed, Low defender speed. (h) Low ball speed, Medium defender speed. (i) Low ball speed, High defender speed.

**Figure S2.**
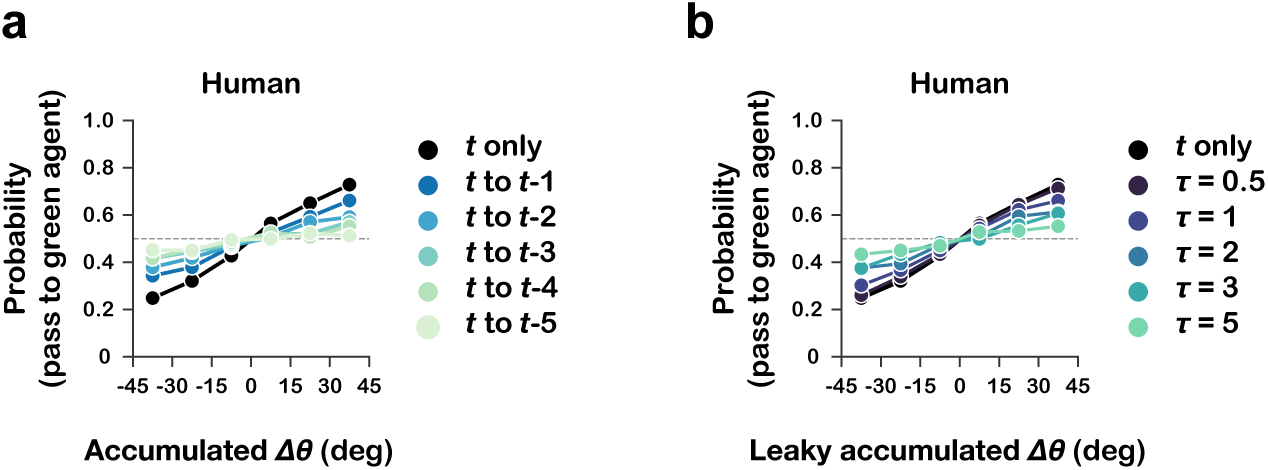
Human passing decisions. (a) Probability of a human passing to the green agent for each Δ*θ*, computed using different accumulation windows. (b) Probability of a human passing to the green agent for each Δ*θ*, computed using different decay weights, with the accumulation window fixed at 5 time steps (500 ms).

**Figure S3.**
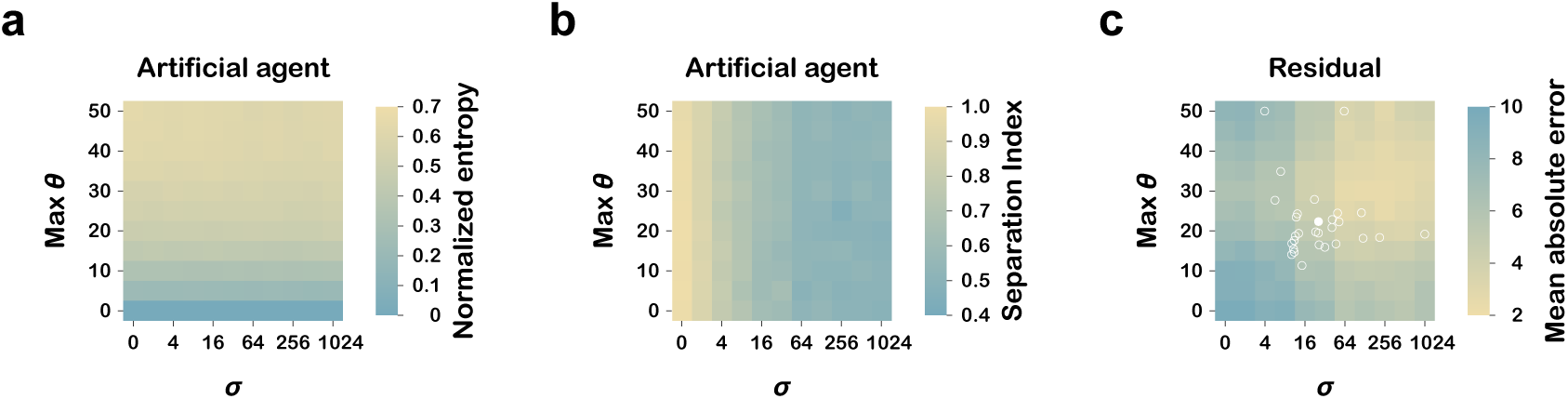
Modeling incorporating the human behavioral characteristics (a) Normalized entropy of the red agent for each condition (combination of Max*θ* and *σ*). For estimating participant entropy, the mean across rows was taken as the value for that Max*θ*. (b) Pass separation of the red agent for each condition. For estimating participant *σ*, the mean across columns was taken as the value for that *σ*. (c) Residuals of performance (mean number of passes) across conditions. Open white circles indicate the estimated Max*θ* –*σ* combinations for individual participants, and the filled white circle indicates their average. The final reproduction of human data was obtained using Max*θ* = 20 and *σ* = 32, which were closest to this average.

**Table S1.** Years of experience in ball sports and non-ball sports for each participant.

| Participant | Ball sports (years) | Non-ball sports (years) |
| --- | --- | --- |
| 1 | 12 | 2 |
| 2 | 11 | 0 |
| 3 | 1 | 12 |
| 4 | 0 | 3 |
| 5 | 10 | 0 |
| 6 | 0 | 10 |
| 7 | 0 | 9 |
| 8 | 4 | 0 |
| 9 | 4 | 4 |
| 10 | 7 | 1 |
| 11 | 12 | 3 |
| 12 | 0 | 11 |
| 13 | 0 | 10 |
| 14 | 0 | 10 |
| 15 | 0 | 3 |
| 16 | 0 | 16 |
| 17 | 0 | 10 |
| 18 | 6 | 3 |
| 19 | 0 | 9 |
| 20 | 0 | 10 |
| 21 | 10 | 0 |
| 22 | 11 | 4 |
| 23 | 0 | 10 |
| 24 | 0 | 5 |
| 25 | 0 | 10 |
| 26 | 0 | 17 |
| 27 | 2 | 17 |
| 28 | 2 | 9 |

**Figure S4.**
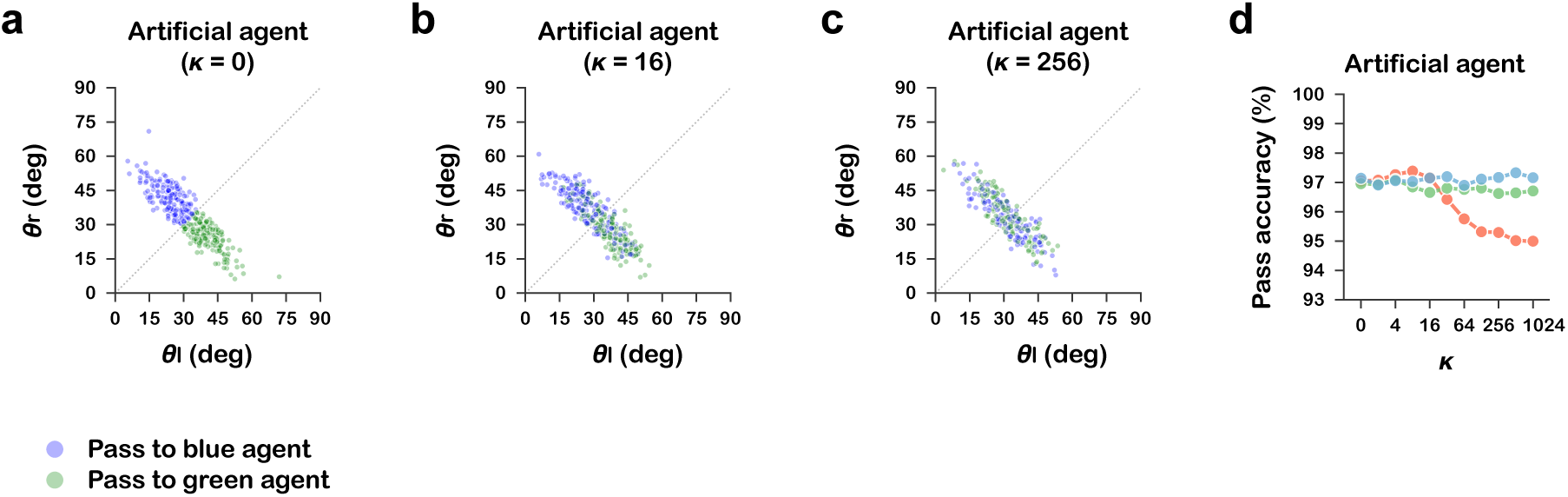
Boltzmann-choice of passing decisions in an artificial agent. (a-c) Passing choices in all-agent teams in which the red agent was assigned different levels of Boltzmann temperature parameter (*κ* = 0, 16, 256). (d) Pass success rates of each agent under different levels of *κ* assigned to the red agent in the all-agent teams. Each line represents one of the three agents (red, green, or blue).

**Figure S5.**
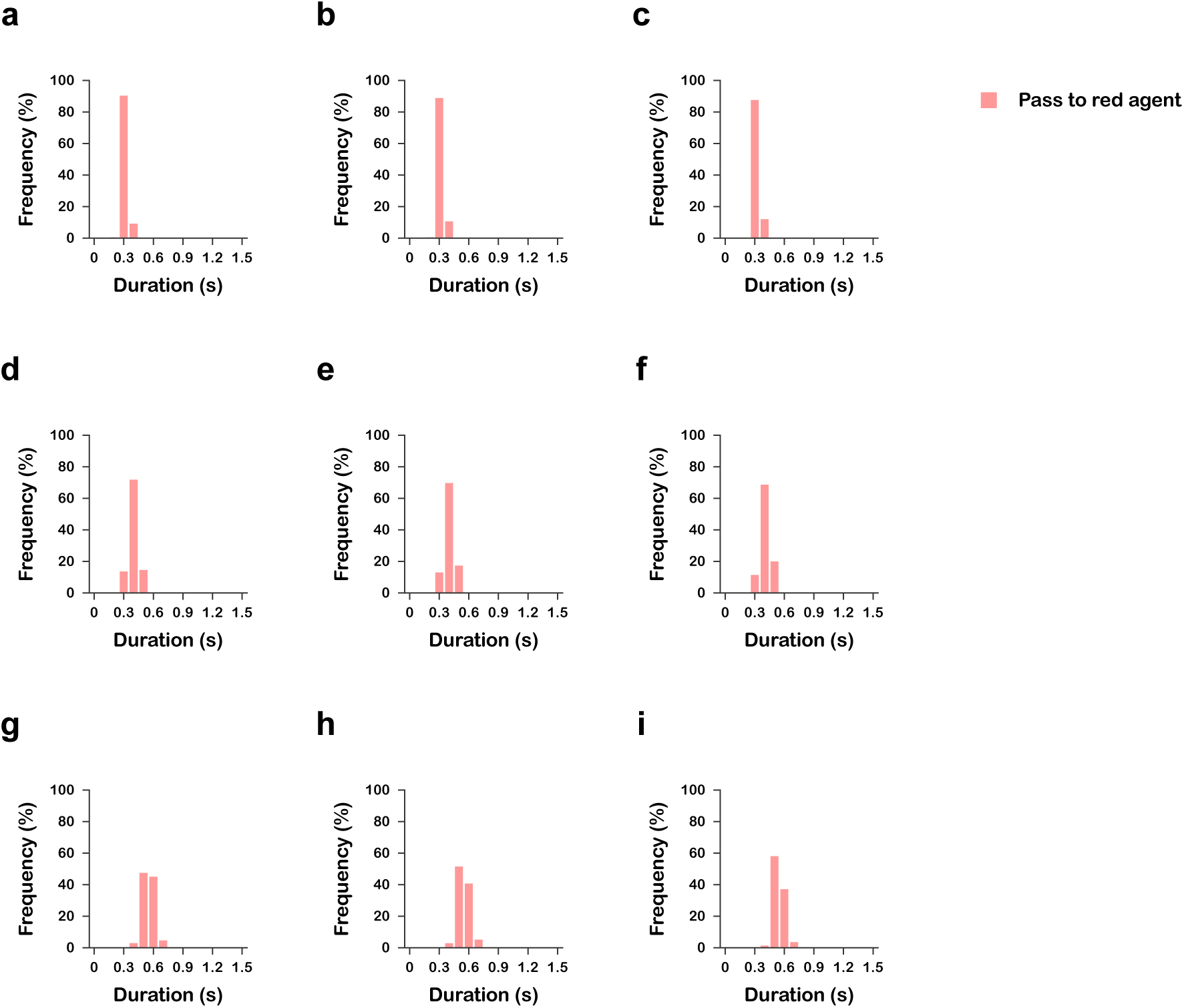
Histograms of the time taken for a pass released by another agent to reach the red agent across the nine experimental conditions. (a) High ball speed, Low defender speed. (b) High ball speed, Medium defender speed. (c) High ball speed, High defender speed. (d) Medium ball speed, Low defender speed. (e) Medium ball speed, Medium defender speed. (f) Medium ball speed, High defender speed. (g) Low ball speed, Low defender speed. (h) Low ball speed, Medium defender speed. (i) Low ball speed, High defender speed.

