## Supplementary figures and images for "Individual bounded rationality destabilizes cooperative dynamics in human–AI groups"

### Supplementary Videos

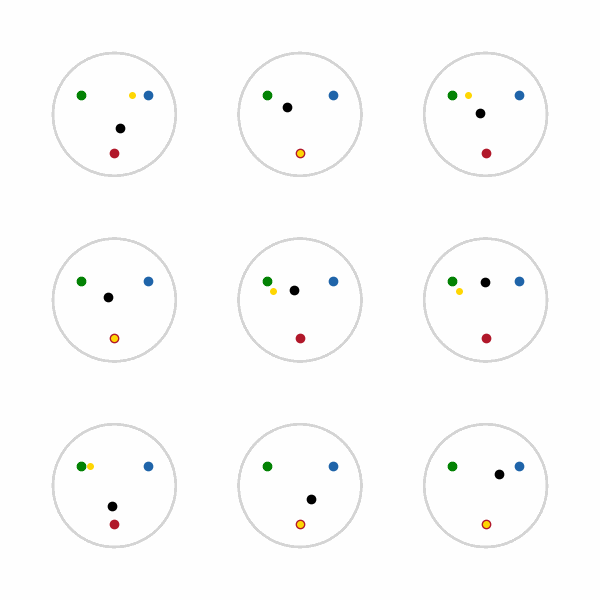
